# VEP-G2P: A Tool for Efficient, Flexible and Scalable Diagnostic Filtering of Genomic Variants

**DOI:** 10.1101/416552

**Authors:** Anja Thormann, Mihail Halachev, William McLaren, David J Moore, Victoria Svinti, Archie Campbell, Shona M Kerr, Sarah Hunt, Malcolm G Dunlop, Matthew E Hurles, Caroline F Wright, Helen V Firth, Fiona Cunningham, David R FitzPatrick

## Abstract

**Purpose:** We aimed to develop an efficient, flexible, scalable and evidence-based approach to sequence-based diagnostic analysis/re-analysis of conditions with very large numbers of different causative genes. We then wished to define the expected rate of plausibly causative variants coming through strict filtering in control in comparison to disease populations to quantify background diagnostic “noise”.

**Methods:** We developed G2P (www.ebi.ac.uk/gene2phenotype) as an online system to facilitate the development, validation, curation and distribution of large-scale, evidence-based datasets for use in diagnostic variant filtering. Each locus-genotype-mechanism-disease-evidence thread (LGMDET) associates an allelic requirement and a mutational consequence at a defined locus with a disease entity and a confidence level and evidence links. We then developed an extension to Ensembl Variant Effect Predictor (VEP), VEP-G2P, which can filter based on G2P other widely used gene panel curation systems. We compared the output of disease-associated and control whole exome sequence (WES) using Developmental Disorders G2P (G2P^DD^; 2044 LGMDETs) and constitutional cancer predisposition G2P (G2P^Cancer^; 128 LGMDETs).

**Results:** We have shown a sensitivity/precision of 97.3%/33% and 81.6%/22.7% for causative *de novo* and inherited variants respectively using VEP-G2P^DD^ in DDD study probands WES. Many of the apparently diagnostic genotypes “missed” are likely false-positive reports with lower minor allele frequencies and more severe predicted consequences being diagnostically-discriminative features.

**Conclusion:** Case:control comparisons using VEP-G2P^DD^ established an observed:expected ratio of 1:30,000 plausibly causative variants in proband WES to ~1:40,000 reportable but presumed-benign variants in controls. At least half the filtered variants in probands represent background “noise”. Supporting phenotypic evidence is, therefore, necessary in genetically-heterogeneous disorders. G2P and VEP-G2P provides a practical approach to optimize disease-specific filtering parameters in diagnostic genetic research.

## Introduction

The analysis of genomic sequence and copy number is now in widespread use as a first-line investigation in the diagnosis of Mendelian disease. In addition to an obvious role in genetic counselling, diagnostic genetic testing can also help avoid invasive procedures (e.g. muscle biopsy in Duchenne and Becker muscular dystrophy^1^) and reduce the length of time required to come to a definitive diagnosis (e.g. leukodystrophies^2^). Such testing has historically been restricted to individuals with distinctive clinical presentations and/or suggestive family histories, which significantly increase the prior probability of specific genetic pathology. However, it is now possible to perform comprehensive analysis of the protein coding region (whole exome sequencing (WES) ^3-5^) or the entirety of the human genome (whole genome sequencing (WGS) ^6,7^) for clinical diagnostic purposes at reasonable cost. Although this represents an exciting opportunity, the number of variants passing any diagnostic filter is strongly correlated with total genomic space sampled. The more genetically heterogenous a disease, the more causal genes are individually implicated and hence the more variants are likely to become diagnostic candidates. It is thus important to develop strategies that can define the impact of increasing the number of variants on false positive and false negative errors in diagnostic assignments; both may result in significant harm through misdiagnosis and missed diagnoses and certainly increase the workload for clinical scientists and clinicians.

The diagnostic filtering of previously unclassified variants is most commonly based on minor allele frequency (MAF) and mutational consequence. The effectiveness of the former has been revolutionized by the availability of data from the Exome Aggregation Consortium (ExAC) ^8^ and the Genome Aggregation Database (gnomAD; http://gnomad.broadinstitute.org). These resources provide access to technically robust variant calls from diverse populations of known providence. There are many different publicly available tools for defining the consequence of an individual variant call^9^. One of the most widely used is the Ensembl Variant Effect Predictor (VEP) ^10^. VEP predicts the effect of each alternative allele on each overlapping transcript for a variant and assigns Sequence Ontology^11^ terms to describe the consequences. It can be run either online or using a locally installed version of the program. VEP exploits the extensive and regularly updated Ensembl datasets to provide the most comprehensive variant annotation possible in coding and non-coding regions. It also allows extensibility through the ‘plugin’ system which allows custom methods to be easily added.

Automated variant annotation and filtering of WES data using the Ensembl VEP has been successfully applied in a genetically heterogeneous disease cohort by the Deciphering Developmental Disorders Study (DDD) ^12,13^. The DDD Study has recruited >13,400 individuals, with previously undiagnosed severe and/or extreme developmental disorders (DD), from the UK and the Republic of Ireland. The principal aim of the project is to define the genetic architecture of DD using trio-based WES analyses as the main analytical tool^14^. Important secondary aims were to identify novel DD loci and develop diagnostic approaches that could be translated into clinical practice. To facilitate this, we developed a database of all known causative DD loci (DDG2P) that was structured to allow facile, very high throughput filtering of variant calls. This dataset has been used in each of the DDD flagship papers^12,13^. The continual updating of DDG2P has been one of the main drivers of the improvement in diagnostic rates through iterative reporting of the same data^15^. The basic architecture and processes used to populate DDG2P^16^ have been adapted to be applicable to any clinical presentation that has a reasonable prior probability of being caused by highly-penetrant genotypes at a defined group of loci.

To expand from DD to other clinical presentations and to create a system that could be maintained and updated by multiple curators, we created the genotype-to-phenotype (G2P) online system with an associated web application to hosts the DDG2P database and any similar datasets.

Here we describe G2P, tailored to address the problem of robust, efficient and flexible prioritization of genotypes identified from NGS data to aid the diagnosis of genetic disease. As part of our G2P system, we have developed a suite of tools and resources: 1. The G2P portal/web-application which is freely available at http://www.ebi.ac.uk/gene2phenotype/ for creation, curation and dissemination of G2P datasets; 2. G2P datasets which formalize collections of locus-genotype mechanism-disease-evidence threads (LGMDET), curated from the literature, and found to be implicated in the cause of a specific disease or clinical presentation; 3. The G2P extension to Ensembl VEP which is freely available at http://www.ebi.ac.uk/gene2phenotype/g2p_vep_plugin(VEP-G2P).VEP-G2P utilizes the allelic requirement information from G2P datasets/panels and leverages allele frequency data from public datasets such as Genome Aggregation Database (gnomAD) together with the mutational consequence annotations from VEP to produce list of potentially causative genotype(s) given an individual’s VCF file as an input. We also present an estimation of false positive and false negative rates associated with its application to WES datasets.

## RESULTS

### G2P data structure, web development and curation interface

The structure of G2P datasets is based on that of the DDG2P diagnostic tool which has been previously described^17^ (**Figure 1A**; Online-only supplemental material). In G2P, each dataset is focused on a disease grouping or defined category of clinical presentation that is of relevance to the clinical diagnosis of Mendelian disease. For a dataset to be publicly released it must be comprehensive, up-to-date and have a plan for future active curation. Each entry in each dataset links a gene or locus, via a disease mechanism, to a disease. A disease mechanism is defined as both an allelic requirement (mode of inheritance, for example biallelic or monoallelic) and a mutation consequence (mode of pathogenicity, for example activating or loss of function). A confidence attribute confirmed, probable or possible is assigned to indicate how likely it is that the gene is implicated in the cause of disease; only confirmed and probable categories are reportable for clinical diagnosis. A fourth category (“both RD and IF”) has been included to highlight for clinical review genotypes that are plausibly associated with both the relevant disease (RD) and another disease that represents an incidental finding (IF). For example, biallelic mutations in *BRCA2* cause a developmental disorder (Fanconi Anaemia) but will also define a cancer predisposition for both parents of the affected individual.

**Figure 1:**
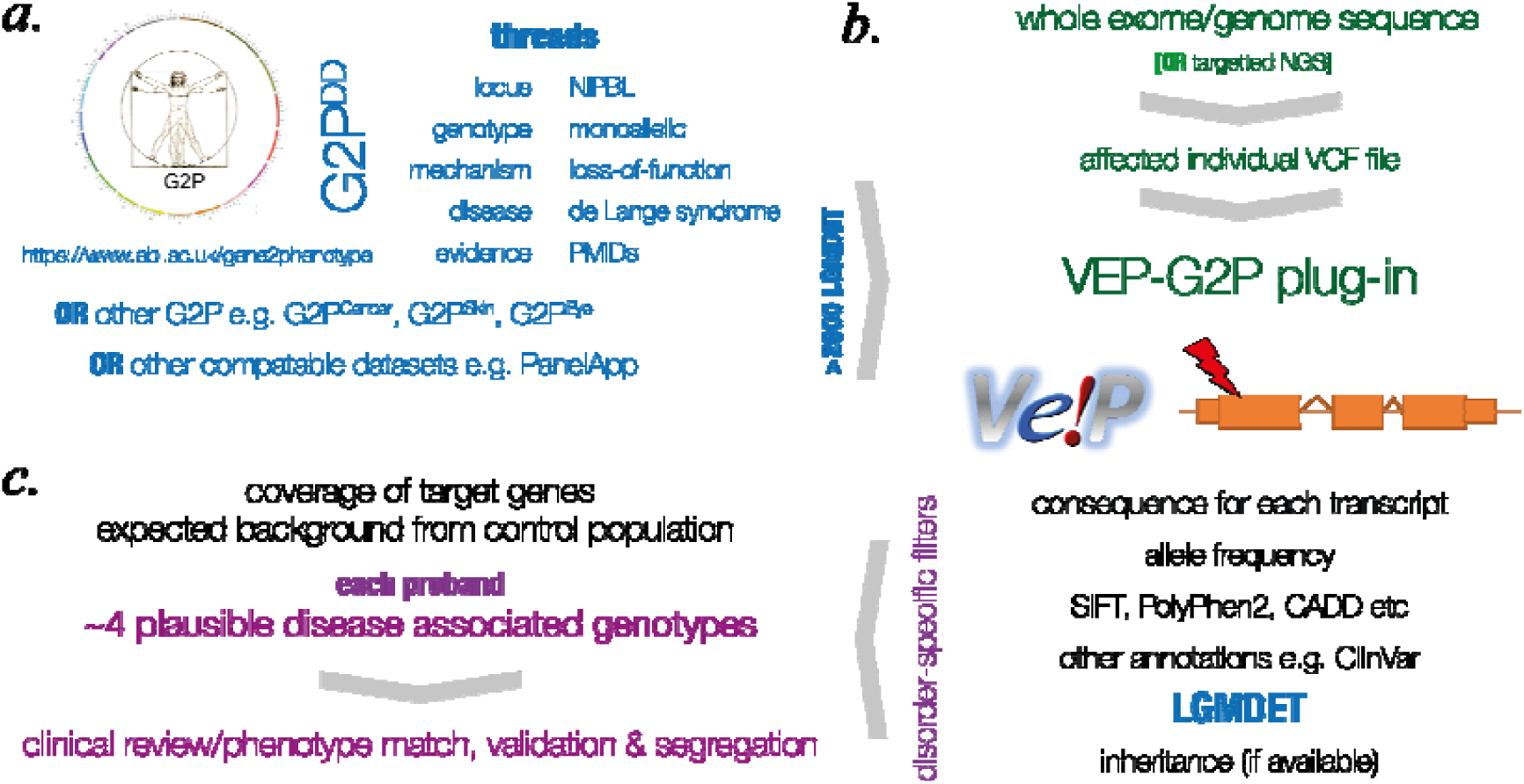
Summary of LGMDET structure and VEP-G2P features. **A.** summarizes the basic logic of the LGMDET approach to genotype classification using an entry for heterozygous, loss-of function variants in *NIPBL* as a cause of Cornelia de Lange Syndrome. The publicly available G2P^DD^ and G2P^Cancer^ data can be searched or downloaded on the website (https://www.ebi.ac.uk/gene2phenotype). Any other compatible dataset, including those developed within PanelApp (https://panelapp.genomicsengland.co.uk), can be used with the VEP-G2P plugin. **B.** The VCF files derived from the next generation sequence data are passed to VEP which uses Ensembl annotation data to compute and annotate the consequence of each variant. The VEP-G2P plugin runs as an additional step of the VEP analysis. It uses the results of VEP’s computations and annotations together with the knowledge from the panel list to filter the variants from the patients input VCF file. The plugin results are returned together with the VEP output file. **C.** The plugin also generates a separate output file which lists the small number of variants and genotypes that pass the variant filtering pipeline implemented in the VEP-G2P plugin– one in HTML format for human use and another in machine readable text format. These genotypes must then be subjected to expert clinical review before any decision regarding causative association with the presenting condition in the affected individual. These variants are at this stage computationally defined and would also normally require validation using a separate technology prior to research or clinical interpretation.

To ensure consistency in development and curation, the rules used to assign confidence, allelic requirement and mutation consequence to entries, are defined and available via the web application in the form of tables (Tables S1-3). We store the links to the publications (via PMID) that provide evidence for that specific gene-disease thread. The locus-genotype-mechanism-disease-evidence link is further characterized by coding the organ specificity and linking to a set of phenotype terms from the Human Phenotype Ontology (HPO) ^17^. These data are all accessible via the G2P web application, which is searchable by gene symbol, disease name or disease ontology term. The released datasets are downloadable as CSV files (https://www.ebi.ac.uk/gene2phenotype/downloads).

### Locus-genotype-mechanism-disease-evidence threads (LGMDET)

Here we present details of the two currently available G2P datasets: G2P^DD^ and G2P^Cancer^ (Table S4 & S5). G2P^DD^ includes LGMDETs associated with clinically significant developmental disorders, i.e.severe and/or extreme disorders that plausibly have their genesis during embryogenesis or early fetal brain developments. This dataset was populated by a combination of clinical knowledge, systematic literature review and genes from existing in-house gene panels by two consultant clinical geneticists (DRF and HVF). We chose to exclude two major groups of developmental disease isolated hearing loss and isolated dental anomalies which are planned to have their own G2P panels and (excepting composite phenotypes) are unlikely to present as undiagnosed developmental disorders. G2P^Cancer^ aims to identify Mendelian cancer susceptibility in individuals affected with cancer or with a strong family history.

The characteristics of the 2044 G2P^DD^ and 123 G2P^Cancer^ reportable (i.e. confirmed, probable, RD&IF) LGMDET entries are summarized in Table 1.

**Table 1:** January 2018 freeze of G2P datasets.

|  | **G2P<sup>DD</sup>** |  | **G2P<sup>Cancer</sup>** |  |
| --- | --- | --- | --- | --- |
|  | **number** | **percent** | **number** | **percent** |
| Reportable* LGMDET | 2044 | 100.0 | 123 | 100.0 |
| Different Reportable Genes | 1517 | NA | 92 | NA |
| **LGMDDET Confidence** |  |  |  |  |
| Confirmed | 1551 | 75.9 | 114 | 92.7 |
| Probable | 403 | 19.7 | 9 | 7.3 |
| Possible** | [257] | NA | [5] | NA |
| RD and IF | 90 | 4.4 | 0 | 0.0 |
| **LGMDDET Allelic Requirement\*\*\*** |  |  |  |  |
| Monoallelic | 701 | 34.3 | 80 | 65.0 |
| Biallelic | 1123 | 54.9 | 38 | 30.9 |
| Digenic | 2 | 0.1 | 0 |  |
| Imprinted | 7 | 0.3 | 0 |  |
| Mitochondrial | 1 | 0.0 | 0 |  |
| Mosaic | 12 | 0.6 | 0 |  |
| Hemizygous | 166 | 8.1 | 2 | 1.6 |
| X-linked dominant and X-linked over-dominance | 32 | 1.6 | 0 |  |
| uncertain | 0 |  | 3 | 2.4 |
| **LGMDDET Mutation Consequence** |  |  |  |  |
| Loss-of-Funtion | 1446 | 70.7 | 107 | 87.0 |
| Activating | 114 | 5.6 | 0 |  |
| Dominant negative | 53 | 2.6 | 0 |  |
| 5' or 3'UTR mutation | 5 | 0.2 | 0 |  |
| Cis-regulatory or promotor mutation | 5 | 0.2 | 0 |  |
| Increased gene dosage | 3 | 0.1 | 0 |  |
| All missense/in-frame | 287 | 14.0 | 6 | 4.9 |
| uncertain | 131 | 6.4 | 10 | 8.1 |

\*Reportable genes are those with a LGMDDET confidence level categorized as probable, confirmed or relevant and incidental. \*\*Possible LGMDETs (see Table S1 for definitions) are not reported in the pipelines used here. \*\*\*An individual gene may have more than one reportable LGMDDET e.g. monoallelic/activating and biallelic/loss-of-function

### VEP-G2P plugin

The VEP-G2P plugin is designed to work with any G2P panel to identify plausibly disease-causing variants from WES or WGS data (VCF files); it enables the facile and flexible integration of allele frequency data in addition to mutation consequence. The default predictions and annotations are invaluable for filtering variants to find those relevant for further analysis based on consequence type and allele frequencies. The VEP-G2P plugin uses the default annotations, the individual’s genotype information and knowledge from the G2P datasets to find genes which have a sufficient number of potentially deleterious variants according to their allelic requirements and are therefore likely disease causing (**Figure 1**; Online-only supplemental material).

### Comparing VEP-G2P outputs from different cohorts

To evaluate the performance of VEP-G2P plugin we analyzed three independent sets of WES data, each of which had undergone extensive prior analysis. The plugin was run using a local VEP installation, and details of the technical aspects of each exome collection are given in Table 2. It should be noted that due to differences in the upstream variant calling pipelines (Table 2; data also processed at different times at different centres), there is a slight excess in the number of filtered and unfiltered indel and SNVs per sample in the DDD cohort compared to the colorectal cancer (CRC) and Generation Scotland (GS) ^18^ cohorts (Table S6). Although it would be possible to realign and recall these datasets to ensure consistency, we chose to proceed without trying to resolve these differences as this is representative of “real life” data available to most research groups involved in clinical diagnostic research.

**Table 2:** Cohort Information and Technical Features WES.

|  | DDD (n=7357)* | CRC (n=517) | GS (n=315) |
| --- | --- | --- | --- |
| Capture kit | Agilent Human All-Exon V3 or V5 Plus<br>with custom ELID C0338371 | Illumina TruSeq Exome Enrichment kit |  |
| Sequencing Platform | Illumina HiSeq | Illumina HiSeq 2000 and 2500 |  |
| Alignment | bwa (0.5.9) | bwa (0.5.9) |  |
| Variant Calling | GATK (3.1.1)<br>Indel realignment, BQSR<br>HaplotypeCaller (run in multisample<br>calling mode using the complete<br>dataset) | GATK (3.4)<br>Indel realignment, BQSR<br>HaplotypeCaller (per sample)<br>GenotypeGVCFs (joint genotyping across all samples on TruSeq<br>regions + 50bp padding) |  |
| Relatedness | After excluding poor quality samples,<br>selected randomly one affected<br>proband per family (using the PED file) | Unrelated | First-degree relatives excluded<br>(based on computed relationship<br>coefficients) |
| Male:Female ratio | 1.36 | 1.11 | 0.73 |
| Median Age | 7.9 yrs | 63 yrs | 52 yrs |

\* DDD details are based on info in the Methods section of “Prevalence and architecture of de novo mutations in developmental disorders” Nature volume 542, pages 433–438 (23 February 2017)

### Discriminative diagnostic indicators

To look for characteristics that may discriminate diagnostic from background genomic variants, we compared the VEP-G2P filtered output of each panel applied to different WES cohorts: The G2P panel with its target disease cohort (i.e. VEP-G2P^DD^ with DDD cohort; VEP-G2P^Cancer^ with CRC cohort), a G2P panel with a discrepant disease cohort (VEP-G2P^DD^ with CRC cohort) and G2P panels with the controls (VEP-G2P^DD^ with GS; VEP-G2P^Cancer^ with GS). The ethical approval and consent procedures governing the recruitment to the DDD Study allow diagnostic analyses only for the identification of pertinent genetic results. It specifically prohibits the intentional identification of “incidental findings”, such as genotypes related to adult-onset cancer susceptibility; for this reason, we did not apply filtering using VEP-G2P^Cancer^ with the DDD cohort.

For VEP-G2P^DD^ the proportion of SNV that survived filtering was 1 in 31.4K in the DDD cohort and 1:44.3K in GS (Table S6; p = 6.56E-13). The rate in the CRC cohort using VEP-G2P^DD^ was similar to the control group at 1 in 40.8K (Table S6; p = 0.16 cf GS). Comparing the results from the DDD cohort with GS controls there is a significant excess of loss-of-function and missense variants for monoallelic and biallelic genes. A higher proportion of the surviving variants in monoallelic genes were missense variants in GS compared to DDD (83.7% cf 64.9%) (**Figure 2A,B**; Tables S7,S8). The missense variants that survived filtering in DDD had a higher proportion with a CADD ^19^ score >30 compared to GS (11.4% cf 7.6%;) (**Figure 2D**). The mean MAF of all missense variants in GS was 1.68x higher than in DDD for monoallelic genes and 1.24x for biallelic genes. For loss-of-function variants the MAF was 2.0-3.5x higher in GS. 115/454 (25.3%) monoallelic DDG2P genes reported had a higher proportion of individuals with variants in GS compared to the DDD WES (Table S13) while 224/454 (49.3%) had no reported variants in GS. The respective proportions for biallelic DDG2P genes are 63/676 (9.3%) and 4/676 (0.6%) (Tables S13).

**Figure 2:**
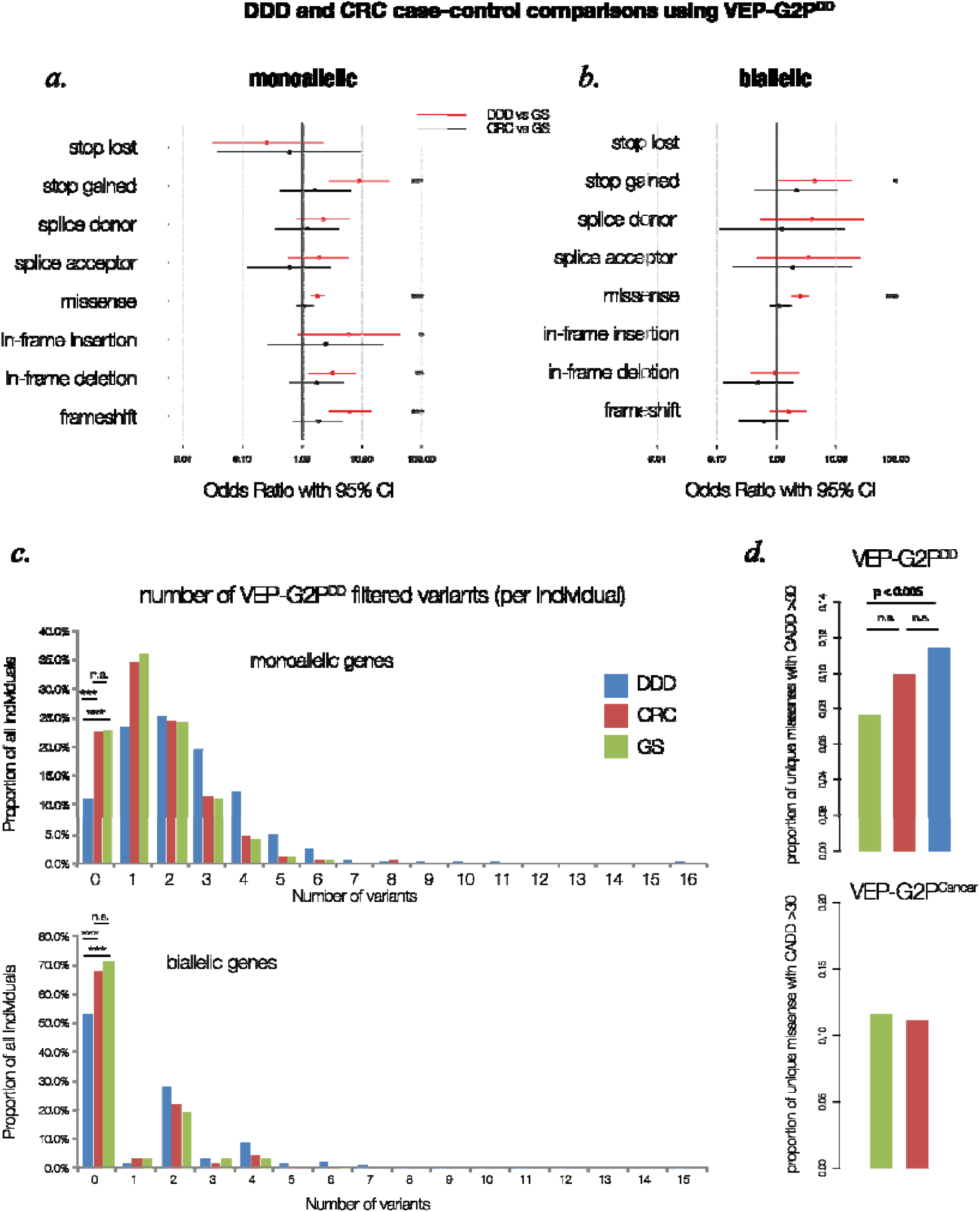
Diagnostically discriminative VEP-G2P disease-specific output. This figure summarizes the VEP-G2P analysis of three independent WES cohorts; DDD (n=7357), CRC (n=517) and GS (n=315) **A.** Odds ratios for samples carrying at least one valid G2P variant (passing the G2P criteria and on a canonical transcript) of specific type in the 454 unique G2P^DD^ monoallelic genes: DDD vs GS (red) and CRC vs GS (black); two-tail Fisher’s Exact Test: ^*^ p-value ≤ 5×10^-2^, ^**^ p-value ≤ 5×10^-3^, ^***^ p-value ≤ 5×10^-6^; considering only missense variants for which both SIFT and PolyPhen agree deleterious/damaging. **B.** Odds ratios for samples carrying at least one valid G2P variant (passing the G2P criteria and on a canonical transcript) of specific type in the 950 unique G2P^DD^ biallelic genes. N/A for stop_lost and inframe_insertion variants no such variants found in the GS cohort, only few found in DDD and CRC (p-value > 5×10^-2^). **C.** Proportion of individuals in the three cohorts (y-axis) carrying a particular number of LOF and missense (regardless of their SIFT/PolyPhen status and CADD score) variants reported by VEP-G2P^DD^ (x-axis). The proportion of DDD individuals for which no VEP-G2PDD hit is found is significantly lower compared to CRC and GS cohorts, both for monoallelic (p-values for two-tail Fisher’s Exact Test comparing number of individuals for which no variants is found to those for which at least one variant is found: DDD vs GS = 7.9e-09, DDD vs CRC= 2.3e-12, CRC vs GS = 0.93) and biallelic genes (DDD vs GS = 1.5e-10, DDD vs CRC = 1.5e-11, CRC vs GS = 0.39). DDD (n=7357 individuals), CRC (n = 517), GS (n = 315). **D.** Proportion of unique missense variants with CADD score > 30 in each of the three cohorts. DDD cohort is significantly enriched for unique missense variants with CADD > 30 in G2P^DD^ genes (top) compared to GS (p-value two-tail Fisher’s Exact Test = 0.005); there is no significant difference between DDD and CRC (p-value = 0.17) and CRC and GS (p-value = 0.16). There is no significant difference for the proportion of unique missense variants with CADD > 30 in the CRC and GS cohorts in G2P^Cancer^ genes (bottom,p-value = 1.0). A relaxation of the uniqueness constraint by accounting for all missense variants found in individuals (regardless of their presence in other individuals in the same cohort) leads to similar results DDD is enriched for missense variants with CADD > 30 in G2P^DD^ genes compared to GS (p-value = 0.01); additionally, CRC cohort also appear enriched for such variants compared to GS (p-value = 0.041); data not plotted.

**Figure 3.**
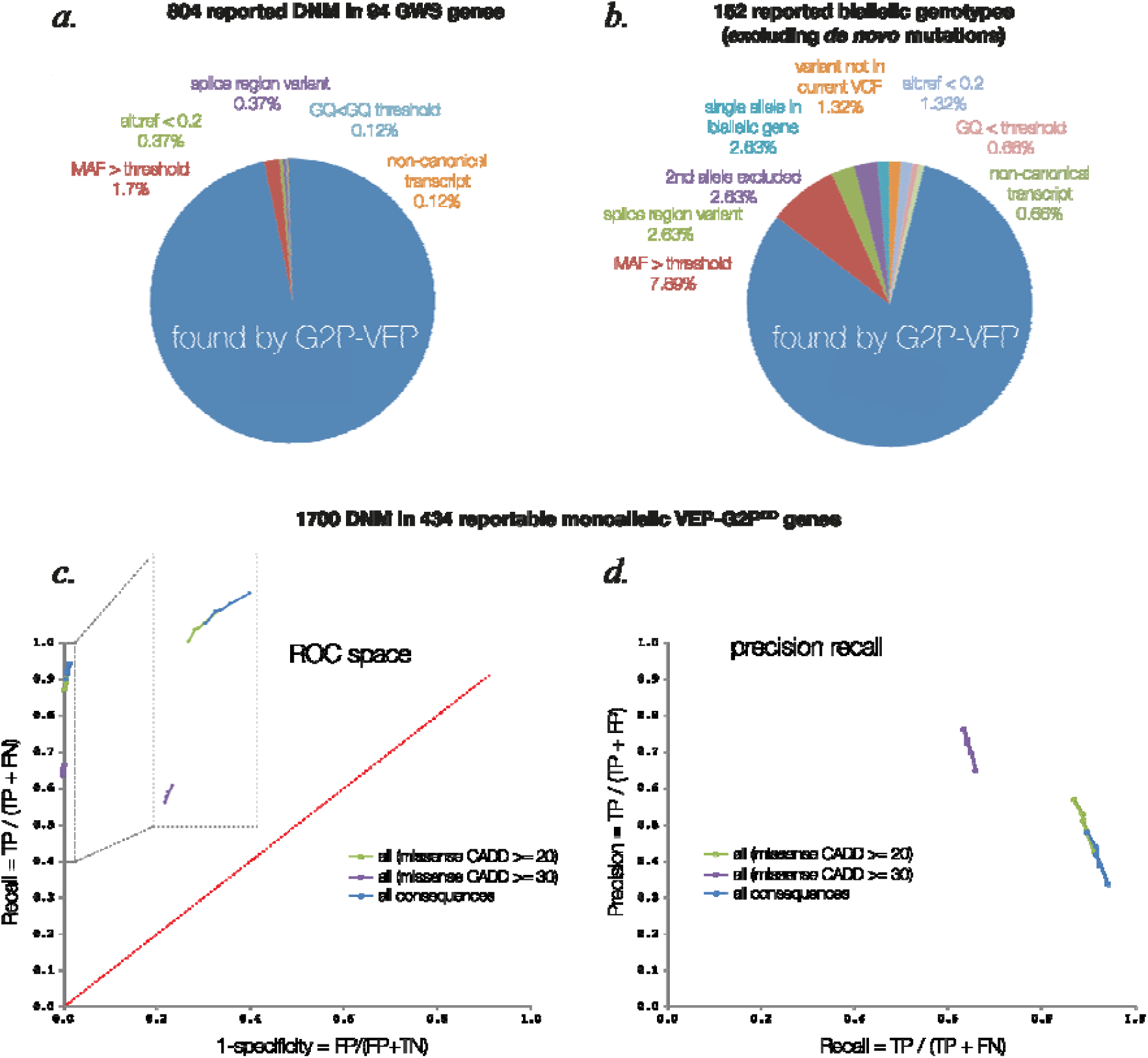
Sensitivity and precision of VEP-G2P Analysis A. Evaluation of G2P accuracy against the set of variants previously identified independently by DDD in 94 genes which reach genome-wide significance (GWS) for *de novo* mutations in the DDD study. Sensitivity = TP / (TP + FN) = 782 / (782 + 22) = 782 / 804 = 97.3%. Precision = TP / (TP + FP) = 782 / (782 + 1522 + 38) = 782 / 2342 = 33.4%. **B.** Evaluation of G2P accuracy against the set of variants previously identified independently by DDD in the first 1133 samples, excluding *de novo* mutations. Sensitivity = TP / (TP + FN) = 124 / (124 + 28) = 124 / 152 = 81.6%. Precision = TP / (TP + FP) = 124 / (124+ 423) = 124 / 547 = 22.7% **C.** The ROC curves for the performance of VEP-G2P^DD^ on the 1700 DDD probands with denovo mutations identified in the 484 DDG2P monoallelic genes by the DDD consortium. We consider as True Negatives (TN) all variants identified in the 484 monoallelic genes which are not reported by VEP-G2P^DD^ (i.e., TP and FP) or missed by VEP-G2P^DD^ but reported in the DDD curated set (i.e., FN). The points on the curves represent varying MAF cut-offs: not seen in any control databases (bottom left), MAF < 1:100000, MAF < 1:50000, MAF < 1:25000, MAF < 1:10000 (top right). To aid visualisation of the data, the region in the top left corner of the ROC space graph has been expanded to scale using the regions bounded by the dashed line rectangles **D.** Evaluation of the consequence type and MAF effects on precision and recall (PR curves) of VEP-G2P^DD^ using the same data analysed for the ROC space in C. The highest precision [0.812, 0.863] is achieved by restricting the analysis only to LOF variants (excluding all missense and inframe insertion/deletion variants); however, this approach leads to the lowest recall [0.425, 0.437]. Conversely, the highest recall is achieved when considering variants of all consequence types [0.897, 0.942] at the cost of decreased precision [0.334, 0.476]. As can be expected, restricting the analysis to consider only missense variants with CADD >= 30 or CADD >= 20 leads to improvements in precision at the cost of decreasing recall. Imposing various MAF thresholds (not seen in any control databases, MAF <= 1:100000, 1:50000, 1:25000, 1:10000; represented as points on each curve, top-to-bottom) affects precision and recall (to a smaller degree).

Using G2P^Cancer^ in the CRC cohort compared to GS revealed no significant enrichment in anyclass of variant (**Figure 2A,B**). However, monoallelic G2P^Cancer^ LGMDET stop-gained variants in GS had a MAF 2.67x higher than those in CRC (Table S11). For biallelic G2P^Cancer^ LGMDET, no variants survived filtering in GS with 9 reported in the CRC WES (Table S12). 25/61 of all genes reported by the VEP-G2P plugin were found in a higher proportion of individuals in GS than CRC with 22/61 being exclusively reported in CRC (Table S14).

Using all variant surviving VEP-G2P^DD^ filtering there was a mean of 3.8 variants per DDD proband (3.59 SNV and 0.19 INDEL; Table S6) compared to 2.14 variants per individual in the GS controls (2.05 SNV and 0.09 INDEL; Table S6). The distribution of the numbers of variants reported per individual is shifted to the right in DDD probands compared to individuals in GS (**Figure 2C**). With a significantly smaller proportion of DDD probands have no variants reported compared to both GS individuals (p = 7.9e-09) and CRC probands (p = 2.3e-12). However, these differences could, at least in part, be systematic and reflect the alignment/variant calling, read depth or targeted pull-down used in each analysis (**Table 2**) rather than any underlying differences in biology of the populations. Analysis of larger control cohorts that have been processed using the same pipelines as the case cohort and the variants jointly called will be required to determine if these differences are real.

### Sensitivity and precision of identifying previously assigned causative variants in DDD cohort

Using data from the first 4293 trio WES in the DDD study the over-representation of plausibly deleterious *de novo* variants in 94 different genes achieved genome-wide significance^12^. There was a total of 804 likely causative *de novo* variants in these 94 genes that were reported to referring clinicians. Proband-only analysis using VEP-G2P with DDG2P LGMDETs (VEP-G2P^DD^) successfully identified 782 (97.3%) of the reported variants. These 782 variants were amongst the 2342 variants that survived filtering, giving a precision of 33.4% and a false positive rate of 66.6%. Of the 22 *de novo* mutations that were missed, the most common reason was that they had a MAF that was higher than the 1:10000 cut-off used in our monoallelic filtering (**Figure 3A**).

To assess the performance of VEP-G2P^DD^ in identifying inherited causative variants, we used the recent comprehensive re-analysis of known diagnoses in the first 1133 trios in DDD^16^ excluding reported *de novo* mutations. This method successfully identified 124 of the 152 known diagnostic inherited variants, giving a sensitivity of 81.6% with a precision of 22.7%. The reasons for “missed” diagnoses were similar to those for de novo mutations (**Figure 3B**).

Receiver operating characteristics (ROC) analysis has proven to be a highly effective method of comparing the performance of diagnostic tests. The most common form of ROC space analysis uses a continuous variable to create a ROC curve the larger the area under this curve (AUC) the “better” the test. We wished to explore how VEP-G2P^DD^ performed in ROC space. We therefore chose to use the set of 1700 de novo mutations which occurred in DDD probands within genes that were monoallelic and reportable in G2P^DDs^. The only continuous variable that is available to us was the MAF and given that our filter cutoff is 1:10,000 and many variants are unique (having no computationally useful MAF) the area of ROC space that can be interrogated using this approach is very small. However we can calculate the lower bound for AUC of 0.964 using the simple approach developed for binary tests^20^ ([sensitivity + specificity]/2) using VEP-G2P default parameters.

It has been noted recently that ROC curve analysis can be misleading when using binary classifiers^21^ and that precision-recall curves may be used in conjunction with ROC curves to provide a more realistic picture of the tests under investigation. The precision-recall plot using the same data as that used in the ROC analysis does indeed show the cost of increasing sensitivity with respect to precision (**Figure 3B**).

## METHODS

### Whole exome sequence (WES) data filtering

The three WES cohorts (Table 2) were screened for poor quality/potentially contaminated samples. For each sample, we computed the number of extreme HET variants (AD/DP < 0.15 or AD/DP > 0.8) and the number of rare HOM variants (ExAC AF < 0.01). Samples with extreme HETs > cohort mean + 3sd or rare HOMs < cohort mean 3sd were excluded from further analyses; there were 159 such samples in DDD and seven in each CRC and GS.

The variants identified for each sample were screened for quality and those with GQ < 13 (95% confidence), DP < 5 (DDD has the lowest average coverage of the three cohorts) or AD/DP < 0.2 were reset to “no-calls”. Furthermore, variants with Mapping quality (MQ) < 13 (95% confidence) in DDD were also reset to “no-calls”; MQ filtering was not possible for the GS cohort (combined VCF, no MQ value available for individual calls). The variants in the cohorts’ VCFs have been decomposed and normalized with VT (v0.5) prior to submission to G2P.

### G2P and VEP-G2P development

Detailed descriptions of G2P and VEP-G2P development and implementation are provided as Online-only supplemental material

## Discussion

DDG2P was developed to identify reportable, plausibly causative genotypes in known developmental disorders in DDD Study probands^13^. Our primary aim was to evaluate LGMDETs the key architectural feature of DDG2P data as a scalable and generalizable approach for diagnostic analysis of clinical presentations in which affected individuals may have one of many different Mendelian disorders. First, we developed gene2phenotype (https://www.ebi.ac.uk/gene2phenotype/) to facilitate the creation and review of LGMDETs in different datasets. To maintain consistency and clarity of purpose in G2P datasets, to date, we have used only two highly-motivated expert clinician curators to develop and maintain each G2P dataset. This approach requires a significant investment of time and effort and is difficult to scale. However, data mining tools (Pubtator, ClinVar etc) are now being incorporated into the online system to minimize the human resource requirements. Additional curation tools will become increasingly important as the diversity of journals reporting novel gene-disease associations continues to widen. Here we present G2P^DD^ and G2P^Cancer^ as first two publicly accessible LGMDET sets.

Our primary aim is also dependent on the ability to implement these LGMDET sets in clinical research diagnostic filtering. For this we chose to develop the VEP-G2P plugin as both G2P and VEP are hosted at EMBL-EBI and VEP is widely used in research and clinical practice. The VEP-G2P plugin identifies and reports genotypes that fulfil LGMDET requirements and MAF filters. G2P filtering aims to report only likely causative genotypes. Reporting genotypes as opposed to lists of plausibly pathogenic variants produces only a small numbers of loci (mean <4), minimizing the time required for review of each case by clinicians and clinical scientists. For specific loci reporting genotypes also masks incidental findings e.g. only homozygous or two different heterozygous (possible compound heterozygous) likely pathogenic variants in *BRCA2* will be reported in DDG2P as a cause of Fanconi anaemia.

The speed and ease of VEP-G2P plugin use has allowed us to assess the expected background output from each G2P dataset against a population ascertained dataset. This required access to WES data from individuals that have not been selected for any specific disease or clinical problem and we have used these individuals as our control group. Here we used Generation Scotland but in the near future, much larger, unselected WES and WGS control datasets will be available from UK Biobank^22^. Our analysis suggests that at least half of the variants surviving the filtering process are likely to be the result of background population genome variation rather than specifically relevant to this disease being analyzed. We consider this to be a very important “sanity check” in genetic diagnostic analysis. For this reason, we have made it very simple for any panel of any size to be converted to be compatible with VEP-G2P plugin. We have been able to implement PanelApp (https://panelapp.genomicsengland.co.uk) gene panel compatibility in VEP-G2P (using a simple flag in the command line) as the data structures are broadly similar to those of G2P. PanelApp currently hosts 231 gene panels focused on specific clinical diseases (e.g. Charcot Marie Tooth syndrome) or on groups of phenotypically-related diseases (e.g. Hereditary Ataxias). These gene panels were mostly initiated using panels in current clinical use with subsequent crowd-sourced curation. No matter what their origin, panels that do not show a clear difference between the output from controls and disease cohort data (using the appropriate MAF and variant consequence filters) require reassessment and/or revision prior to implementation for clinical or research use. Such analysis will be particularly useful to identify genes with a very restricted repertoire of disease-associated variants and a high background of rare high-impact variants. Such loci may be better analyzed using a variant “whitelist” approach.

Determining the “diagnostically-useful completeness” of any panel in any curation system is a major challenge; requiring balancing all possible associations of a set of comparable genotypes with the clinical presentation against the confidence that the association is causative rather than coincidental. We have found both the statistical genomics analysis (identifying loci achieving genome-wide significance under different genetic models) and clinician case updates within DDD very helpful for DDG2P curation. However, it will be important to establish robust methods to quantitate this feature in any clinical presentation.

Family-trio WES data are hugely valuable for determining the *de novo* status of variants in monoallelic genes, as well as the phase of potential compound heterozygous variants in biallelic genes. In the absence of trio data, there is a particular problem associated with accurate calling of genotypes in ultra-rare biallelic disorders as evidenced by the expected high rate of false positives which is the result of an inability to differentiate variants *in cis* and *in trans* using VEP-G2P^DD^ for proband-only analyses, where it is not possible to determine the phase of most variants detected within a single gene. This will be helped by longer read technology and deeper, more comprehensive data on background genetic variation in human populations. It is interesting that a significant proportion of the “missed” diagnoses in our *de novo* analysis were due to variants previously being assigned as causative which, on current analyses, show implausibly high MAF values.

Finally, we would like to emphasize that the VEP-G2P plugin should be considered a system for experts and it is not designed for use by laboratories or clinical services who do not have competence and experience in a multi-disciplinary approach to the diagnosis of rare genetic disease involving both scientists and clinicians. Casual use of this system could result in misdiagnosis and subsequent significant mismanagement of ‘affected’ individuals.

## Acknowledgements

Ensembl receives majority funding from Wellcome (grant numbers WT095908, WT098051, WT108749/Z/15/Z). This project has received funding from Wellcome (WT200990/Z/16/Z) and the European Molecular Biology Laboratory. This project has received funding from the European Union’s Horizon 2020 research and innovation programme under grant agreement n° 634143 (MedBioinformatics). The DDD study presents independent research commissioned by the Health Innovation Challenge Fund [grant number HICF-1009-003], a parallel funding partnership between Wellcome and the Department of Health, and the Wellcome Sanger Institute [grant number WT098051]. The views expressed in this publication are those of the author(s) and not necessarily those of Wellcome or the Department of Health. The study has UK Research Ethics Committee approval (10/H0305/83, granted by the Cambridge South REC, and GEN/284/12 granted by the Republic of Ireland REC). The research team acknowledges the support of the National Institute for Health Research, through the Comprehensive Clinical Research Network. This study makes use of DECIPHER (http://decipher.sanger.ac.uk), which is funded by the Wellcome. HF is supported by the Wellcome Trust [award 200990/Z/16/Z] ‘Designing, developing and delivering integrated foundations for genomic medicine’. The views expressed in this publication are those of the author(s) and not necessarily those of the Wellcome Trust or the Department of Health. The research team acknowledges the support of the National Institute for Health Research, through the Comprehensive Clinical Research Network. Funding for UK10K was provided by the Wellcome Trust under award WT091310. DRF funded as part of the MRC Human Genetics Unit grant to the University of Edinburgh. MH is supported by an IGMM Translational Science Award. Generation Scotland received core support from the Chief Scientist Office of the Scottish Government Health Directorates [CZD/16/6] and the Scottish Funding Council [HR03006]. Genotyping of the GS:SFHS samples was carried out by the Genetics Core Laboratory at the Wellcome Trust Clinical Research Facility, Edinburgh, Scotland and was funded by the Medical Research Council UK and the Wellcome Trust (Wellcome Trust Strategic Award “STratifying Resilience and Depression Longitudinally” (STRADL) Reference 104036/Z/14/Z).

## Online resources

Gene2Phenotype:http://www.ebi.ac.uk/gene2phenotype/

VEP-G2P https://www.ebi.ac.uk/gene2phenotype/g2p_vep_plugin

PanelApp https://panelapp.genomicsengland.co.uk

## References

1. Darras BT, Miller DT, Urion DK. Dystrophinopathies. 1993;GeneReviews.

2. Parikh S, Bernard G, Leventer RJ, et al. A clinical approach to the diagnosis of patients with leukodystrophies and genetic leukoencephelopathies. Mol Genet Metab. 2015;114(4):501–515.

3. Biesecker LG. Exome sequencing makes medical genomics a reality. Nat Genet. 2010;42(1):13–14.

4. Choi M, Scholl UI, Ji W, et al. Genetic diagnosis by whole exome capture and massively parallel DNA sequencing. Proc Natl Acad Sci U S A. 2009;106(45):19096–19101.

5. Ng SB, Buckingham KJ, Lee C, et al. Exome sequencing identifies the cause of a mendelian disorder. Nat Genet. 2010;42(1):30–35.

6. Gilissen C, Hehir-Kwa JY, Thung DT, et al. Genome sequencing identifies major causes of severe intellectual disability. Nature. 2014;511(7509):344– 347.

7. Lupski JR, Reid JG, Gonzaga-Jauregui C, et al. Whole-genome sequencing in a patient with Charcot-Marie-Tooth neuropathy. N Engl J Med. 2010;362(13):1181–1191.

8. Ruderfer DM, Hamamsy T, Lek M, et al. Patterns of genic intolerance of rare copy number variation in 59,898 human exomes. Nat Genet. 2016;48(10):1107–1111.

9. Pabinger S, Dander A, Fischer M, et al. A survey of tools for variant analysis of next-generation genome sequencing data. Brief Bioinform. 2014;15(2):256– 278.

10. McLaren W, Gil L, Hunt SE, et al. The Ensembl Variant Effect Predictor. Genome Biol. 2016;17(1):122.

11. Eilbeck K, Lewis SE, Mungall CJ, et al. The Sequence Ontology: a tool for the unification of genome annotations. Genome Biol. 2005;6(5):R44.

12. Study DDD. Prevalence and architecture of de novo mutations in developmental disorders. Nature. 2017;542(7642):433–438.

13. Study DDD. Large-scale discovery of novel genetic causes of developmental disorders. Nature. 2015;519(7542):223–228.

14. Firth HV, Wright CF, Ddd S. The Deciphering Developmental Disorders (DDD) study. Dev Med Child Neurol. 2011;53(8):702–703.

15. Wright CF, McRae JF, Clayton S, et al. Making new genetic diagnoses with old data: iterative reanalysis and reporting from genome-wide data in 1,133 families with developmental disorders. Genet Med. 2018.

16. Wright CF, Fitzgerald TW, Jones WD, et al. Genetic diagnosis of developmental disorders in the DDD study: a scalable analysis of genome-wide research data. Lancet. 2015;385(9975):1305–1314.

17. Klhler S, Vasilevsky NA, Engelstad M, et al. The Human Phenotype Ontology in 2017. Nucleic Acids Res. 2017;45(D1):D865–D876.

18. Smith BH, Campbell A, Linksted P, et al. Cohort Profile: Generation Scotland: Scottish Family Health Study (GS:SFHS). The study, its participants and their potential for genetic research on health and illness. Int J Epidemiol. 2013;42(3):689–700.

19. Kircher M, Witten DM, Jain P, OÕRoak BJ, Cooper GM, Shendure J. A general framework for estimating the relative pathogenicity of human genetic variants. Nat Genet. 2014;46(3):310–315.

20. Cantor SB, Kattan MW. Determining the area under the ROC curve for a binary diagnostic test. Med Decis Making. 2000;20(4):468–470.

21. Saito T, Rehmsmeier M. The precision-recall plot is more informative than the ROC plot when evaluating binary classifiers on imbalanced datasets. PLoS One. 2015;10(3):e0118432.

22. Sudlow C, Gallacher J, Allen N, et al. UK biobank: an open access resource for identifying the causes of a wide range of complex diseases of middle and old age. PLoS Med. 2015;12(3):e1001779.

